# A biosensor-based phage display selection method for automated, high-throughput Nanobody discovery

**DOI:** 10.1101/2024.06.26.600780

**Authors:** Phebe De Keyser, Valentina Kalichuk, Thomas Zögg, Alexandre Wohlkönig, Stephan Schenck, Janine Brunner, Els Pardon, Jan Steyaert

## Abstract

Biopanning methods to select target-specific Nanobodies^®^ (Nbs) involve presenting the antigen, immobilized on plastic plates or magnetic beads, to Nb libraries displayed on phage. Most routines are operator-dependent, labor-intensive and often material- and time-consuming. Here we validate an improved panning strategy that uses biosensors to present the antigen to phage-displayed Nbs in a well. The use of automated Octet biolayer interferometry sensors (Sartorius) enables high throughput and precise control over each step. By playing with association and dissociation times and buffer composition, one can efficiently decrease the background of aspecific and low-affinity Nbs, reducing the rounds of panning needed for the enrichment of high-affinity binders. Octet panning also enables the use of unpurified target proteins and unpurified phage from a bacterial culture supernatant. Additionally, downscaling to a 384-well format significantly reduces the amount of protein required. Moreover, enrichment of binders can be quantified by monitoring phage binding to the target by interferometry, omitting additional phage titration steps. Routinely, up to three rounds of Octet panning can be performed in only five days to deliver target-specific binders, ready for screening and characterization using the same Octet instrument.

**Graphical abstract:** 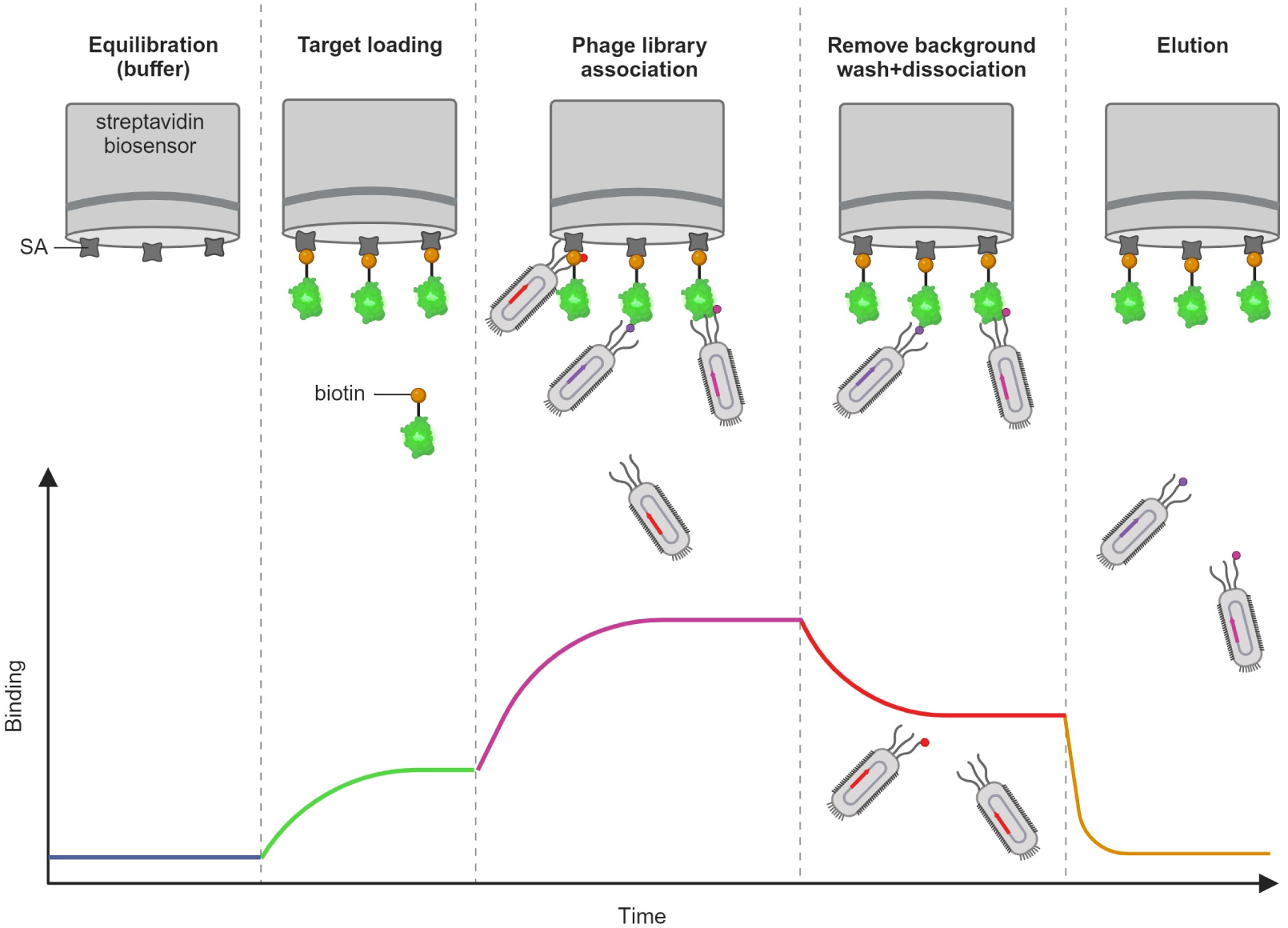

## Introduction

The production of recombinant proteins in microbial systems has revolutionized biochemistry by enabling protein characterization and the development of protein-based therapeutics (Rosano and Ceccarelli, 2014). Nevertheless, many proteins, especially eukaryotic proteins, integral membrane proteins and multi-protein complexes are still challenging targets to study because they are difficult to produce, purify or characterize (Elmore and Coaker, 2011; Saccardo et al., 2016).

Nanobodies^®^ (Nbs), the small (around 15 kDa) variable domains of heavy-chain only antibodies that naturally occur in camelids (Hamers-Casterman et al., 1993), are valuable research tools to study such ‘difficult’ proteins. For example, Nbs can be used as affinity reagents to purify proteins (Stevens et al., 2024) or co-expressed with membrane proteins to act as stabilizing chaperones that enhance their expression (Claes et al., 2016). Nbs have also been applied for the structural characterization of several important protein classes (Laverty et al., 2019; Rasmussen et al., 2011b, 2011a). Conformation-specific Nbs can be used to freeze dynamic proteins into a single conformation that can be analyzed by cryogenic electron microscopy (Uchański et al., 2020) or to lock targets in druggable conformations for drug discovery (Pardon et al., 2018). Nbs can also be introduced as intracellular antibodies (intrabodies) to visualize a protein and study the link between its structural dynamics and cellular behavior (Dmitriev et al., 2016; Irannejad et al., 2013; Rothbauer et al., 2006). Additionally, their small size and high solubility, stability and affinity cause a growing interest in the therapeutic potential of Nbs (Jovčevska and Muyldermans, 2019).

Most pipelines for the discovery of target-specific Nbs involve the immunization of a llama with the target (protein) to create an immune library, but naïve or synthetic libraries have also been screened successfully. Target-specific binders from these immune libraries-displayed on phage, yeast, bacteria or mammalian cells-are next enriched using biopanning or similar selection methods (Boder and Wittrup, 1997; Francisco et al., 1993; Ho et al., 2006; Smith, 1985; Winter et al., 1994). In conventional biopanning, the target is immobilized in the wells of a plastic microtiter plate and presented to phage-displayed Nbs (Pardon et al., 2014). Next, the plate is manually washed repeatedly to remove aspecific phage. Phage that are specifically bound to the target are eluted typically by changing the pH or by using trypsin to proteolytically release the phage from the target. Since the elution step also releases aspecific phage, all eluted phage, enriched for target-specific binders, are amplified to use as input for a following round of selection. The whole procedure is time-consuming and labor-intensive and the washing steps, which define the stringency of the panning, are operator-dependent. An additional disadvantage of presenting the target on a plate is the lack of control over target orientation and functionality. To overcome these shortcomings, alternative panning methods were developed, for example by presenting the target on a column (Noppe et al., 2009) to increase washing efficiency and eliminate the need for multiple selection rounds. Presenting the target on magnetic beads or surface plasmon resonance biosensors enables automated manipulations using a KingFisher™ Flex (Kumaran et al., 2012) (ThermoScientific, Waltham, MA, USA) or WHITE FOx™ (Donck et al., n.d.) (FOx BIOSYSTEMS, Diepenbeek, Belgium) respectively. The FOx biosensor-based method also allows to monitor the immobilization of the target and the binding of phage to it. However, all these methods still take weeks to perform multiple rounds, include time-consuming steps like phage purification and titration, and use considerable quantities of the purified target.

Individual clones from the output of a biopanning are traditionally screened for binding using an enzyme-linked immunosorbent assay (ELISA). However, the ELISA signal is highly influenced by the Nb expression yield and does not allow for an accurate ranking of the Nbs based on binding kinetics. Therefore, Ylera et al. developed a method to do quantitative dissociation rate-based ranking of antibodies directly from lysates on an Octet^®^ (Sartorius, Göttingen, Germany) (Ylera et al., 2013). This robust biolayer interferometry (BLI)-based instrument dips a biosensor in consecutive wells of a plate and monitors biomolecule interactions that cause a shift Δλ (nm) in the optical interference pattern on the biosensor surface.

Here we validate a novel, fast and versatile biopanning method that presents the target on such a biosensor to phage-displayed Nbs in a well. The use of automated Octet biosensors ensures robustness, reproducibility and precise control over each step. Since most Nbs have association rate constants k_on_ of 10^5^-10^6^ M^−1^ s^−1^ while their dissociation rate constants k_off_ vary more (Muyldermans, 2013), most affinity can be gained by selecting Nbs that dissociate slowly from the target. Building on this principle, Octet panning can easily be adapted to select either a high variety of low-affinity binders or a lower variety of high-affinity binders. By directly monitoring phage binding to the immobilized target by interferometry, separate phage titration can be omitted. Time-consuming purification steps can be reduced as well, since this method enables the use of (nanogram amounts of) unpurified target and unpurified phage from a bacterial culture supernatant. As a result, three consecutive rounds of panning can be performed in only five days. To further increase the throughput or for performing comparative panning experiments, 8 (Octet Red 96/Octet R8), 16 (Octet Red 384/Octet RH16) or even 64 (Octet RH96) pannings can be done in parallel while keeping tight control over all volumes, incubation times and buffer compositions, followed by screening and characterization of the most promising binders using the same instrument.

## Material and methods

### Simulations of the binding kinetics of the antigen-Nb interaction on an Octet sensor

The number of Nbs bound on a target-functionalized streptavidin (SA) Octet sensor was simulated during association and dissociation using MATLAB R2021b. Association was simulated for 10 min in a 200 µL well containing 1 pM of binding Nbs and dissociation for a 200 µL well without binders. The association rate constant was set to be 4 × 10^5^ M^−1^ s^−1^ and the dissociation rate constant was varied from 0.04 to 40 × 10^−3^ s^−1^. The sensor was set to have an area of 0.28 mm^2^ with max 10^9^ target binding sites. A 1:1 binding model was used for the simulations (Appendix A).

### Selection of GFP-specific Nbs using Octet

A llama was immunized with Green Fluorescent Protein (GFP) and an immune library with a size of 8.8 × 10^7^ independent clones was prepared and purified as described in Pardon et al. (Pardon et al., 2014) using pHEN4 as the M13 phagemid vector and TG1 *Escherichia coli* (*E. coli*) as the host. The purified phage library was stored at a concentration of ~6 × 10^12^ phage/mL, estimated using absorbance at 260 nm (Lee et al., 2007).

Pannings were performed using an Octet Red 96 in Flat Bottom 96-well plates (Greiner, cat. no. 655076) using SA sensors (Sartorius, cat. no. 18-5019). SA sensors were hydrated 10 min in PBSTB (PBS pH 7.4, 0.005 % Tween 20, 0.01 % BSA) outside of the instrument. For each step of the selection method, a separate well of the 96-well plate was filled with the appropriate solution. Each time, a well volume of 200 µL was used, the sample plate was shaken at 1000 rpm and kept at 30 °C during the experiment. GFP (Appendix B (Scholz et al., 2000)) was biotinylated using EZ-Link™ NHS-PEG4-Biotin (ThermoFisher, cat. no. A39259) according to the manufacturer’s instructions. Biotinylated GFP was loaded onto sensors until the binding signal increased ~1 nm. After a 30 s wash and 30 s re-equilibration in PBSTB, GFP-sensors were incubated for 10 min with 1 nM (6 × 10^11^ phage/mL) of the purified phage library. After a 3 min wash in PBSTB, sensors were transferred stepwise to wells containing PBSTB allowing phage to dissociate from the sensors first for 10 min and next for 90 min. Phage still bound on GFP were eluted by transferring the sensors to a well with 200 mM glycine at pH 2.3 for 1 min. Phage eluted with acid were neutralized with 1 M Tris-HCl pH 9.5 to a final concentration of 120 mM Tris-HCl (Trizma Sigma-Aldrich, cat. no. T6066). As the negative control, this panning was mirrored with a sensor that was loaded with an irrelevant biotinylated protein of 15 kDa.

### Screening of GFP-specific Nbs by BLI on Octet

To recover phage from a well, 50 µL was incubated with 1 mL exponentially grown TG1 cells for 30 min at 37 °C and supplemented with 5 mL TY/ampicillin/glucose to grow overnight at 37 °C, shaking at 200 rpm. Single colonies from the overnight cultures were grown, induced to express Nbs and lysed by freezing and thawing, based on the protocol in Pardon et al. (Pardon et al., 2014). For screening, these lysates were diluted fourfold in PBSTB. For association, Octet SA sensors loaded with GFP were incubated for 10 min in Nb containing lysate. Dissociation was next monitored by plunging these sensors for 10 min in PBSTB. As a positive control, we used a lysate containing Nb207. Nb207 is a pM-affinity GFP binder discovered previously from the same GFP immune library via conventional panning (Steyaert et al., 2021). As a negative control, a lysate containing an irrelevant Nb was used.

Association and dissociation sensorgrams were analyzed in MATLAB, first smoothing the data and subtracting a reference curve (PBS instead of lysate) before aligning all curves at the start of association. By convention, Nbs were classified as antigen-specific binders when their signal was > 0.06 nm higher compared to the negative control at the end of association. Antigen-specific binders were ranked based on the stability of the complex, measured as the time required for 50 % dissociation. A weak binder has a half-life shorter than 1 min. A medium binder has a half-life between 1 and 10 min, whereas a strong binder has a half-life exceeding 10 min. All binders were sequenced and grouped into the same family when the complementarity-determining region 3 (CDR3) has the same length and more than 80 % sequence identity (Pardon et al., 2014). For expression, Nbs were recloned into pMESy4 (GenBank KF415192), expressed with a His6-tag in WK6 *E. coli* cells and purified via Immobilized Metal Chelate Affinity Chromatography (IMAC) (Pardon et al., 2014). Full kinetic characterizations were done on Octet Red 384 following Sartorius’ guidelines. A similar kinetic characterization of Nb207 was done in an Octet R8 with evaporation cover (Sartorius, cat. no. 18-5132).

### Selection of mCherry-specific Nbs using Octet

A llama was immunized with 1 µg mCherry Sku (Uniprot X5DSL3) resulting in an immune library with a size of 3.4 × 10^8^ clones. For Octet pannings, biotinylated mCherry was loaded onto sensors until the signal increased ~1 nm. An Octet panning was performed as described above, incubating mCherry-sensors with the purified phage library. An extra 30 s wash step was added before elution to avoid transfer of free phage in liquid sticking around the sensor. For the second round of panning, 50 µL of eluted phage were recovered overnight, rescued with helper phage, amplified and prepared as described in Pardon et al. (Pardon et al., 2014). mCherry-specific Nbs were identified from the second round by screening on Octet as described above, using mCherry on the sensor.

### Selection of FBA1-specific Nbs using Octet

A llama was immunized with the soluble yeast proteome of *Saccharomyces cerevisiae* strain EBY100 (ATCC® MYA-4941™), resulting in an immune library with a size of 2.5 × 10^8^ clones. To prepare the proteome, EBY100 cells were grown in YPD, harvested in mid-log phase, resuspended in PBS supplemented with 50 μg/mL DNAse I and EDTA-free protease inhibitors (Roche), and lysed with a French press. The lysate was centrifuged and the supernatant containing all soluble proteins was filtered with a 0.22 µm filter.

Pannings were performed as described above, using biotinylated Nb207 on the sensors to capture separate yeast proteins that were fused to GFP. Yeast 1,6-bisphosphate aldolase (FBA1, Systematic Name YKL060C) was expressed as a GFP-fused protein in an engineered yeast strain from the Yeast GFP fusion collection (Huh et al., 2003). FBA1-GFP was captured on the sensor from an overnight culture lysed with Y-PER™ Plus (ThermoScientific) diluted fourfold in PBSTB. Lysate from the EBY100 reference strain was used to prepare control sensors.

After each round of panning, 50 µL of eluted phage were rescued and prepared for a following round. The third round was screened as described. SA sensors were loaded with biotinylated Nb207 to capture FBA1-GFP from the yeast lysate or GFP alone from an *E. coli* lysate.

### Comparison of different methods to elute phage bound to antigen-loaded Octet sensors

Three elution methods to release bound phage from an antigen-loaded sensor were compared: 1 min in 200 mM glycine pH 2.3, 20 min in 200 mM glycine pH 2.3 or 20 min in 0.25 mg/mL trypsin. GFP-biotin was loaded onto SA sensors until the signal increased ~1 nm. After 30 s wash and 30 s re-equilibration in PBSTB, the sensor was incubated for 10 min with 2 nM phage displaying Nb207, washed for 1 min and eluted under the different test conditions. Phage in the glycine elutions were neutralized, while trypsin activity was inhibited with AEBSF. The number of eluted phage was counted by infecting TG1 cells and pipetting dilutions on LB/ampicillin/glucose plates (Pardon et al., 2014).

### Monitoring the binding of phage to antigen-loaded Octet sensors

To monitor the binding of phage to antigen-loaded sensors, we equilibrated SA sensors, loaded GFP-biotin until ~1 nm, washed for 30 s, re-equilibrated 30 s and incubated with Nb207 (control) *versus* Nb207-phage for 10 min at different concentrations. Association curves were also obtained for different dilutions of the supernatant (can be stored for at least six weeks at −20 °C) of an *E. coli* culture infected with Nb207-phage.

### Octet selection of GFP-specific Nbs starting from unpurified phage

To select GFP-specific Nbs starting from an unpurified phage library, a 20 mL culture of the GFP immune library was infected with helper phage and grown overnight at 37 °C in a shaking 50 mL tube. The culture supernatant was collected by spinning down the TG1 cells for 15 min at 3200 rcf. GFP-biotin was loaded onto SA sensors until saturation by incubating them for 1 min at 20 µg/mL. After 30 s wash and 30 s re-equilibration in PBSTB, one sensor was incubated for 10 min with the culture supernatant containing the phage. As a positive control, another sensor was dipped in 1 nM of the phage, now purified from the same culture supernatant following the standard protocol. After a 3 min wash in PBSTB, phage were dissociated from the sensors for 100 min in PBSTB. After an extra 30 s wash step, remaining phage were eluted in 200 mM glycine at pH 2.3 for 1 min. As negative controls, purified and unpurified phage displaying an irrelevant Nb were panned in parallel with GFP-loaded sensors.

After this first round of panning, 50 µL of eluted phage were rescued. The first and second rounds were performed in 200 µL wells of a 96-well plate (Octet R8). The second round was also performed in 80 µL wells of a Flat Bottom 384-well plate (Greiner, cat. no. 781076) and 40 µL wells of a Tilted Bottom 384-well plate (Greiner, cat. no. 18-5080) on an Octet Red 384.

Starting from the unpurified phage library, we also panned for high-affinity GFP binders by including two long dissociation steps of 400 min, bringing the total dissociation time to 15 h. To avoid evaporation, this experiment was carried out at 15 °C and an evaporation cover was used. All pannings were screened as described above.

### Selection of IgSF8-specific Nbs using Octet starting from unpurified phage

A llama was immunized with mouse brain extracts enriched in synaptic vesicles (Takamori et al., 2006) resulting in an immune library with a size of 6.7 × 10^8^ clones. To prepare the target antigen, HEK293T cells were grown in a 6-well culture plate (ThermoFisher, cat. no. 140675) in DMEM with 10 % FBS (Life Technologies) and transfected with EGFP-fused mouse brain membrane protein IgSF8 (Uniprot G3UYZ1) using X-treme GENE 9 DNA Transfection Reagent (Sigma Aldrich, cat. no. 6365787001). Twenty-four hours after transfection, cells were lysed for 2 h at 4 °C on a rocking platform with 0.5 mL per well of 20 mM HEPES pH 7.5, 150 mM NaCl, 5 % glycerol, 1 % DDM/0.1 % CHS, DNAse I and protease inhibitors (Roche) and spun for 30 min at 20800 rcf at 4 °C. A negative control was obtained following the same procedure without adding DNA to the Transfection Reagent. To measure in-gel fluorescence, the supernatants were run in a 4-15 % mini-protean TGX gel (Bio-Rad, cat. no. 4561084) next to PageRuler™ prestained protein ladder (ThermoFisher, cat. no. 26616). An image of the gel was made on the white and blue trays of a Gel Doc EZ (Bio-Rad).

Pannings were performed on an Octet R8 using SA sensors saturated with Nb207 by incubating them for 1 min in 10 µg/mL Nb207-biotin. Next, the sensors were saturated by plunging them in 200 µL lysate of IgSF8-EGFP transfected cells or lysate of control cells. After 30 s wash and 30 s re-equilibration, sensors were incubated for 2 h with the unpurified phage library supplemented with protease inhibitors. After two 1 min wash steps, remaining phage were eluted in 200 mM glycine at pH 2.3 for 1 min. All steps were done at 15 °C with 20 mM HEPES pH 7.5, 150 mM NaCl, 5 % glycerol, 0.3 % DDM/0.03 % CHS as assay buffer. For screening the second round, Nb207 was loaded on the sensor to capture IgSF8-EGFP or GFP alone. Binders were grouped in sequence families and two representatives of different families were purified via IMAC. Both Nbs and an irrelevant one were coupled to 50 µL NHS-agarose beads (Cytiva, cat. no. 17090601) for a 1 h immunoprecipitation of IgSF8-EGFP from 250 µL lysate. After washing the beads three times with assay buffer, IgSF8-EGFP in-gel fluorescence was detected as described above.

### Semi-throughput selection of yeast protein-specific Nbs using Octet

Parallel pannings were performed to select binders against five GFP-fused yeast proteins from an immune library raised against the soluble yeast proteome, starting from unpurified phage. Each antigen was loaded on a different sensor: 1,6-bisphosphate aldolase (FBA1, YKL060C), pyruvate decarboxylase isozyme (PDC1, YLR044C), HSP70 type SSA1 (SSA1, YAL005C), phosphogluco-isomerase (PGI1, YBR196C) and 14-3-3 protein (BMH1, YER177W). Pannings were performed essentially as described for the low-throughput panning of FBA1-GFP, now with 10 min dissociation. After a first round, 100 µL of eluted phage were recovered by infecting TG1s and growing for 5 h, rescued and amplified overnight to proceed to a second round the next morning. The second round was screened for binding to the GFP-fused targets or GFP alone as described.

## Results

### Proof of concept experiments

To estimate the margins of panning experiments on Octet, we first simulated the association and dissociation kinetics of phage-displayed Nbs on a target-functionalized SA sensor as shown in Figure 1. From these calculations, it appears that mostly binders with an affinity *K_D_* smaller than 10 nM are still bound on the sensor after 10 min of passive dissociation. After 100 min mostly binders with a *K_D_* smaller than 1 nM are still bound. Mostly binders with a *K_D_* smaller than 100 pM are still bound after 15 h (not shown in the graph). Dissociation times are independent of the phage concentration and reassociation rarely occurs because the phage concentration is neglectable when diluted in a fresh well containing only buffer.

**Figure 1:**
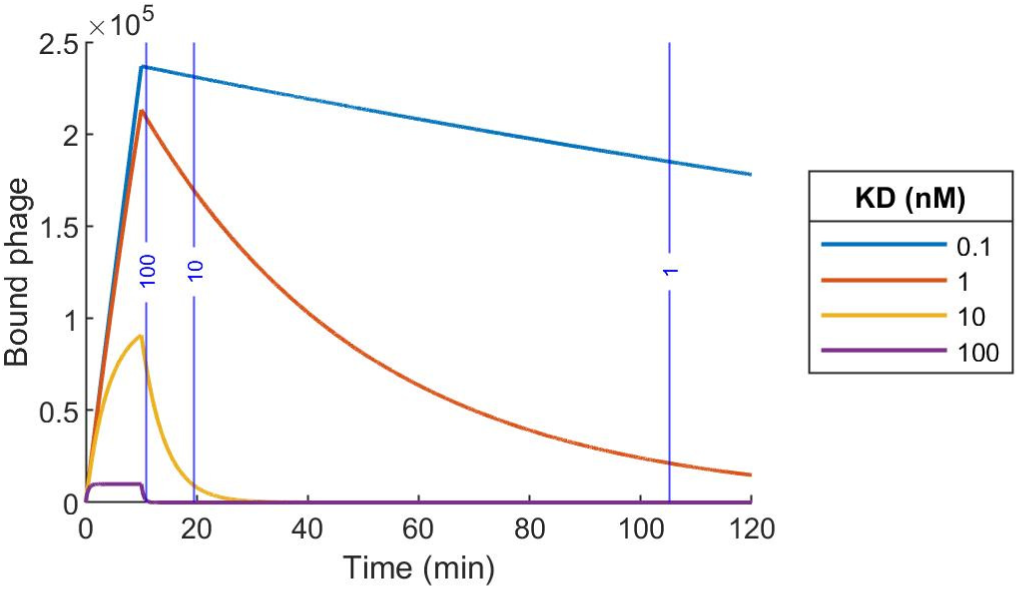
Simulated association and dissociation of phage-displayed Nbs in Octet. The number of phage-displayed Nbs bound to a target-functionalized streptavidin sensor is shown for different affinities of the Nb for the target. Vertical lines indicate the times to dissociate 90 % of phage for different affinities. A 1:1 Langmuir binding model was used. Input parameters: association rate constant = 4 10^5^ M^−1^ s^−1^, sensor surface = 0.28 mm^2^, # binding sites = 10^9^, association: 200 µL, 1 pM of binders, dissociation: 200 µL, 0 pM of binders.

Octet allows to timely control the incubation of an immobilized antigen with a phage library followed by the dissociation of phage in buffer. In a first experiment we used Octet to present GFP that was loaded on a streptavidin functionalized sensor to a Nb library displayed on phage derived from a llama that was immunized with GFP (Pardon et al., 2014). The sensor was incubated in 1 nM of the purified phage library, allowing association for 10 min. After a short wash step, phage were allowed to dissociate for 10 min and 90 min in consecutive wells containing only buffer. Finally, phage still bound on the sensor were eluted at low pH. Phage were recovered from relevant wells, Nbs were expressed and their lysates were screened in the Octet and classified (Table 1, experiment 1). To test the procedure more extensively, a second identical selection was performed (Table 1, experiment 2). All 50 binders were sequenced and grouped in 16 sequence families (Appendix C.1). The family of Nb207, previously discovered from the same library by conventional panning, reappeared but all other families were new. Two representative clones, Nb394 (from the first panning) and Nb395 (from the repetition), were purified (1 mg and 4 mg from 1 L culture, Appendix B) and characterized by BLI on Octet (Appendix C.2). For comparison, Nb207 was characterized as well (Appendix C.3). Similar to Nb207, Nb394 and Nb395 show pM affinity for GFP (Appendix C.4).

**Table 1:** Classification of screened clones of Octet panning with GFP, mCherry and FBA1-GFP. In a first experiment we used Octet to present GFP that was loaded on a streptavidin functionalized sensor to a Nb library displayed on phage derived from a llama that was immunized with GFP. Classification of the GFP-specific binders resulting from this panning are shown: weak binders (half-life < 1 min), medium binders (1 min ≤ half-life < 10 min) and strong binders (10 min ≤ half-life). An identical panning was done in experiment 2, with a phage library prepared from the same immune library. Similar pannings were performed on mCherry (experiment 3) and FBA1-GFP (experiment 4).

| Experiment | 1 | 1 | 1 | 2 | 3 | 4 |
| --- | --- | --- | --- | --- | --- | --- |
| Target | GFP | GFP | GFP | GFP | mCherry | FBA1-GFP |
| Round | 1 | 1 | 1 | 1 | 2 | 3 |
| Phage purified (Y/N) | Y | Y | Y | Y | Y | Y |
| Panning step | 10 min dissociation | 100 min dissociation | Elution | Elution | Elution | Elution |
| Clones screened | 16 | 16 | 13 | 21 | 20 | 45 |
| Target-specific binders (%) | 50 | 75 | 77 | 95 | 100 | 100 |
| Weak binders (%) | 6 | 6 | 8 | 0 | 65 | 80 |
| Medium binders (%) | 31 | 31 | 15 | 14 | 35 | 20 |
| Strong binders (%) | 13 | 38 | 54 | 81 | 0 | 0 |

A similar panning was performed on an immune library raised against purified mCherry (Table 1, experiment 3). To further test the power of our approach we also selected binders against yeast protein FBA1 fused to GFP from an immune library that was raised against a lysate containing all soluble yeast proteins (Table 1, experiment 4).

### Operational improvement of Octet panning

First, we compared different elution methods to recover phage bound onto the sensor (Appendix C.5). Remarkably, trypsin cleavage does not elute considerably more phage than passive dissociation in PBS. Acid elution in glycine at pH 2.3 is almost as efficient for 1 min as for 20 min. Second, we compared selections from purified phage *versus* unpurified phage contained in the supernatant of the bacterial culture (Appendix C.6) and observed minor differences. In general, less phage were eluted starting from the culture supernatant, but the enrichment, calculated as the number of phage eluted in round 2 relative to round 1 was comparable (Appendix C.7). Last, we found that the binding of phage causes changes in the BLI signal that vary depending on the conditions. For example, we prepared Nb207-phage and compared the binding of (1) this phage contained in the culture supernatant, (2) the purified phage and (3) purified Nb207 to a sensor loaded with GFP (Appendix C.8). For the same concentration, purified phage gave a bigger signal change than Nb alone. Remarkably, the purified phage particles caused a decrease in signal whereas the same phage contained in the supernatant caused positive signals. When using immune libraries, binding of phage to the sensor only caused measurable signals in round 2 (Figure 2 and Figure 3). Downscaling the volumes caused a decrease of the amplitude of the association and dissociation sensorgrams (insert in Figure 3).

**Figure 2:**
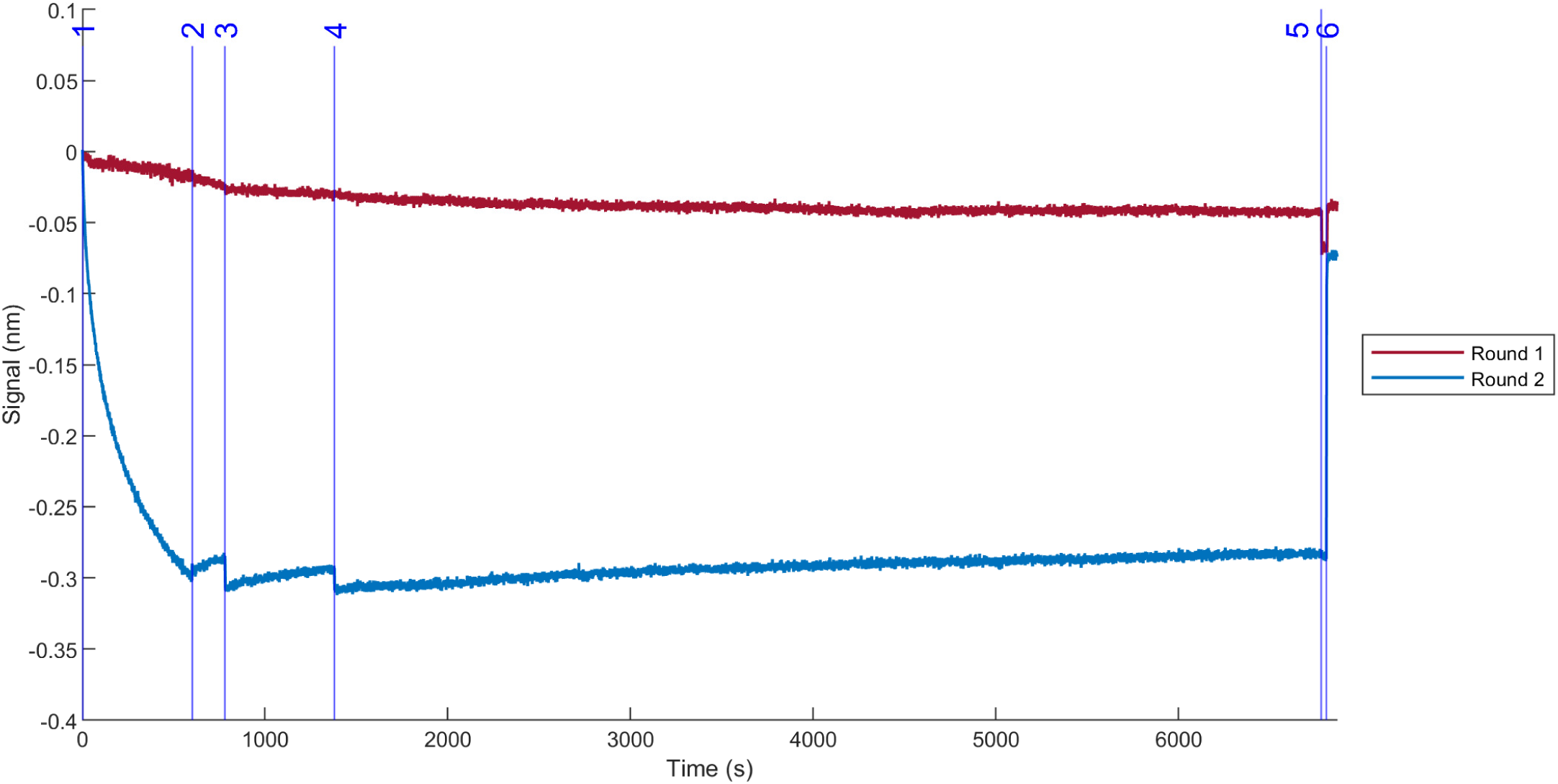
Sensorgrams during Octet panning on GFP with purified phage for round 1 and round 2. Biotinylated GFP was loaded on a streptavidin sensor and the following steps were done in PBS/Tween/BSA: (1) 10 min association with 1 nM (6 10^11^ phage/mL) of the purified GFP phage library (2) 3 min wash, (3) 10 min dissociation, (4) 90 min dissociation, (5) 0.5 min wash, (6) 1 min elution with 200 mM glycine pH 2.3. Eluted phage were rescued and amplified for a second round. As a negative control, a sensor was incubated with irrelevant Nb-phage. The panning curves were analyzed in MATLAB, first smoothing the data and subtracting the irrelevant Nb-phage curves before aligning all curves at the start of association.

**Figure 3:**
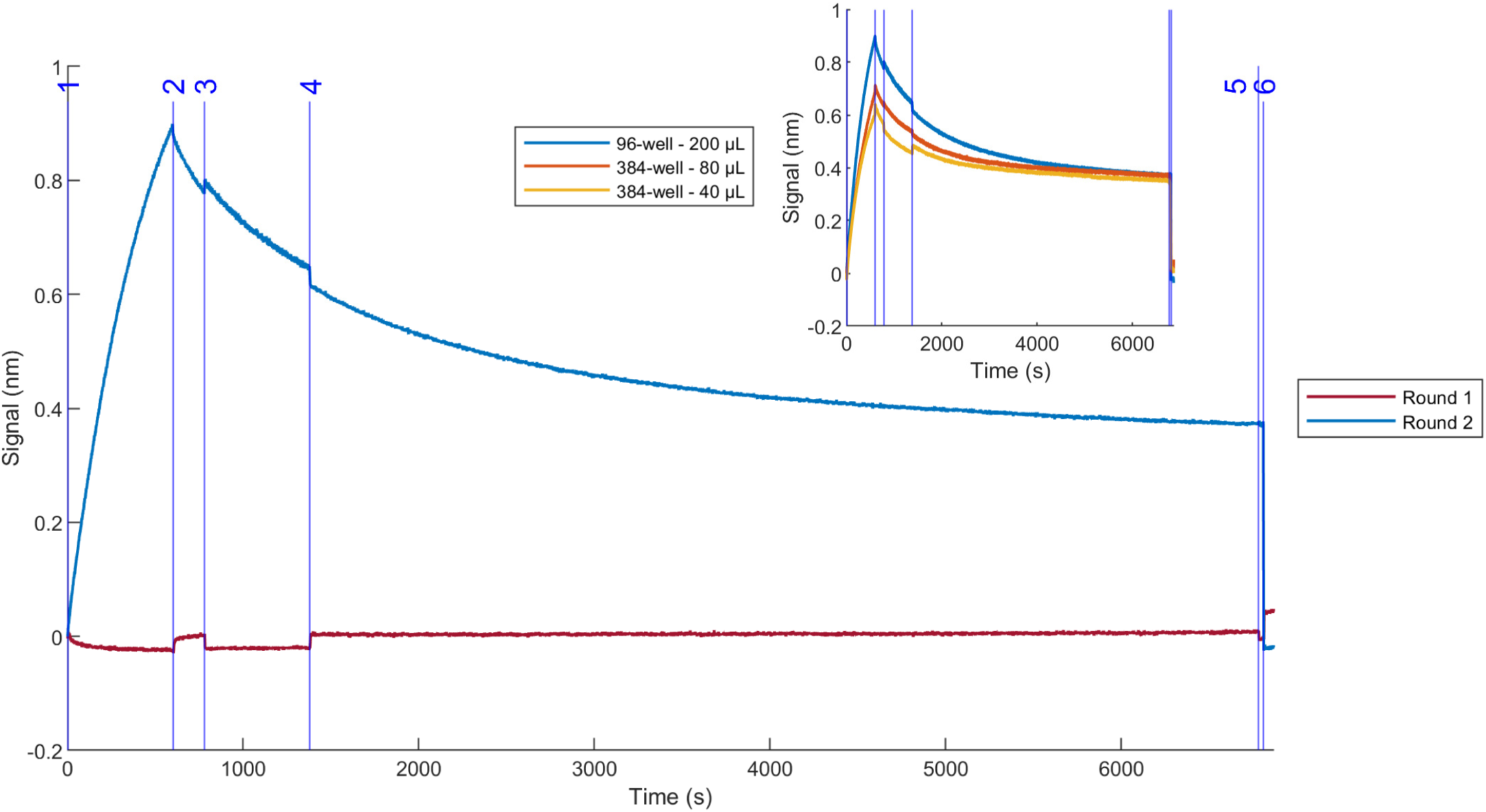
Sensorgrams during Octet panning on GFP with unpurified phage for round 1 and round 2. Biotinylated GFP was loaded on a streptavidin sensor and the following steps were done in PBS/Tween/BSA: (1) 10 min association with culture supernatant containing the GFP phage library, (2) 3 min wash, (3) 10 min dissociation, (4) 90 min dissociation, (5) 0.5 min wash, (6) 1 min elution with 200 mM glycine pH 2.3. Eluted phage were rescued and amplified for a second round. As a negative control, a sensor was incubated with an irrelevant Nb-phage culture supernatant. The panning curves were analyzed in MATLAB, first smoothing the data and subtracting the irrelevant Nb-phage curves before aligning all curves at the start of association. Round 2 was repeated in 80 µL wells of a flat bottom 384-well plate and in 40 µL wells of a tilted bottom 384-well plate (insert).

### Selection of diverse and high-affinity binders by Octet panning

First, we compared binders recovered from phage that were retained more than 15 h on the immobilized antigen to phage that were recovered after 10 min (Appendix C.9). As expected, the diversity of binders decreased significantly for phage that release slower. However, the proportion of very strong binders increased from 0 % after 10 min dissociation to 37 % for phage that were recovered after 15 h.

The Octet panning on membrane protein IgSF8 fused to EGFP also yielded 24 of the 45 screened clones that were specific binders for IgSF8 of which 44 % were classified medium and 9 % strong. Expression of full-length IgSF8-EGFP was confirmed on a gel and we showed for two IgSF8-specific Nbs (Nb776, Nb777) that they can be produced as recombinant proteins (4 mg and 7 mg from 1 L culture, Appendix B) that immune-precipitate IgSF8 (Appendix C.10).

We also used semi-throughput Octet pannings to select specific Nbs against FBA1-GFP and four other GFP-fused yeast proteins from the same immune library derived from a llama that was immunized with the soluble fraction of a yeast lysate. Increasing Octet signals observed during association in round 1 and 2 showed a high enrichment of binders for PDC1-GFP (Figure 4.b) and PGI1-GFP (Figure 4.d). Further analysis yielded low- to high-affinity binders against three out of five antigens (Appendix C.11).

**Figure 4:**
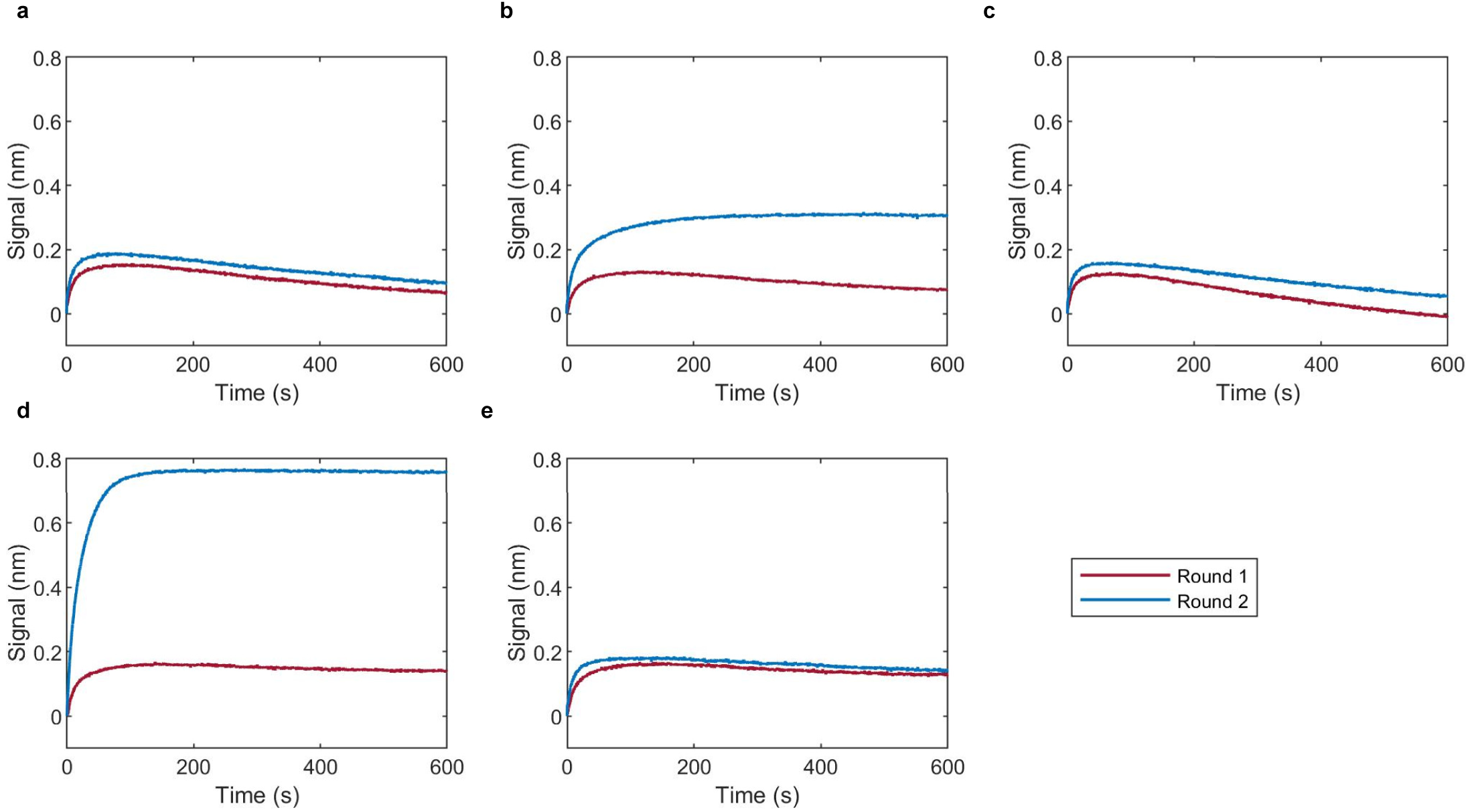
Association curves during Octet panning on GFP-fused yeast proteins with unpurified phage for round 1 and round 2. Biotinylated Nb207 was loaded on streptavidin sensors to capture **a**. FBA1-GFP, **b**. PDC1-GFP, **c**. SSA1-GFP, **d**. PGI1-GFP, **e**. BMH1-GFP from yeast lysate in PBS/Tween/BSA. The curves show the subsequent 10 min association with unpurified phage culture supernatant. After a 10 min dissociation and 1 min elution with 200 mM glycine pH 2.3, 100 µL of eluted phage were rescued and amplified overnight to proceed to a second round the next morning. The curves were aligned at the start of association in MATLAB.

## Discussion and conclusions

### Rationale of the Octet panning method

Most current panning techniques use harsh conditions to elute all phage bound to an immobilized antigen, irrespective of their binding kinetics. Simulations of the association and dissociation kinetics of standard Nbs to common antigens however indicate that most Nbs with an affinity above 10 nM dissociate spontaneously in 10 min whereas pM-affinity binders stick to the antigen for at least 100 min (Figure 1). This simple observation pushed us to develop new strategies to perform pannings in a time-controlled manner that exploit these kinetic features for the selection of high-affinity binders. Immobilizing target proteins on Octet sensors allowed us to incubate these targets with solutions containing phage and elution buffers in parallel experiments under time- and temperature-controlled conditions. With newer models of Octet like R8, RH16 and RH96, cooling and the use of an evaporation cover make it possible to perform experiments overnight.

### Focused discovery of pM-affinity binders by Octet panning

As a proof of concept, we performed a single-round Octet panning on GFP in duplicate, where phage released by passive dissociation in the first 10 min and the following 90 min were collected in consecutive wells containing only buffer and next eluted in a well containing acid. Table 1 shows that the portion of target-specific binders, a measure for enrichment, increases over the consecutive steps of the first panning. Remarkably, this simple protocol yielded a high proportion of binders (77 % in the first panning, 95 % in the repetition), most of these with high affinity (54 %, 81 %). This performance can be attributed to well-controlled washing and dissociation steps. Additionally, the low number of potential binding sites on an Octet sensor compared to ELISA plates used in conventional pannings, may reduce the background and increase the correlation between the abundance of clones and their affinity (Glanville et al., 2015).

As the model predicted, the fraction of strong binders also increases with longer dissociation times. Phage eluted by acid after 100 min dissociation, contained most of the strong binders. The affinities of these binders were not determined case-by-case, but considering a half-life of more than 10 min and assuming a k_on_ of 4 × 10^5^ M^−1^ s^−1^, these binders have an affinity of less than 3 nM, similar to what the model predicted. Additionally, we observed that Nbs that belong to the same sequence family tend to exhibit similar affinities (Appendix C.1). For example, family 1 is a family of medium-affinity binders, while family 2 is a family of high-affinity binders, though most clones in this family showed heterogeneous binding (Apiyo, 2022). Family 3 is a family of high-affinity binders and contains Nb207 (Steyaert et al., 2021), a pM-affinity GFP binder previously discovered from the same library by conventional panning. Two new families containing pM-affinity binders were also characterized whereof Nb394 (from the first panning) and Nb395 (from the repetition) are the representatives. In the past, we have often seen that two repetitive pannings performed under seemingly identical conditions result in the selection of unrelated binders. Off-rate screening on unpurified samples predicted pM-affinities for both Nbs and this was confirmed by a full kinetic characterization using purified Nbs. These experiments show the potential of Octet pannings for the focused discovery of pM binders.

### Low backgrounds persist, also for unpurified targets

Similar to the pannings on GFP, Octet pannings on other antigens, including mCherry and FBA1, also showed remarkably low backgrounds. The panning on FBA1 was performed by capturing GFP-tagged FBA1 from a crude yeast lysate with GFP-specific Nb207 that was immobilized on the sensor. It thus appears that Octet panning can be used for (parallel) high-throughput selections starting from unpurified targets that can be captured on the sensor via a (fusion-)tag. This is especially useful for membrane proteins, which are often fused to GFP to monitor their heterologous expression (Rana et al., 2018).

### Well-timed dissociation can be combined with acid elution of remaining phage

Octet pannings can be used to timely recover phage that remained bound to the immobilized target during passive dissociation. We tested different elution methods (Appendix C.5) and showed that incubating the sensors for 1 min in glycine pH 2.3 is sufficient to elute most remaining (highest-affinity) phage. Upon elution, the BLI signals return to baseline levels instantaneously as seen in Figure 2 and Figure 3.

### Faster Octet pannings using unpurified phage

We performed successful Octet pannings with unpurified phage contained in the supernatant of a bacterial culture (Appendix C.6), omitting the labor-intensive phage purification step before each round of selection. The concentration of phage in the supernatant of an overnight culture is in the order of 10^10^ phage/mL. This is 10 times lower than the titers of purified phage as they are used in conventional pannings. Accordingly, significantly less phage were eluted in round 1 and round 2 if we compare two similar pannings using purified and unpurified phage (Appendix C.7). In both experiments, the backgrounds were so low that it was impossible to calculate conventional enrichment factors as the ratio of phage eluted from the immunized target divided by the number of phage eluted from an irrelevant target. Instead, we looked at the number of acid-eluted phage from round 2 compared to round 1 (both rounds performed under identical conditions) where we eluted 200 times more phage from the immobilized target in both experiments.

### Phage binding to the sensor can be measured at pM concentrations

Association curves of Nb207 to immobilized GFP show that the Octet can monitor this interaction up to ~0.4 nM Nb207 (Appendix C.8.a). Since their big size compared to Nbs causes a bigger change in the sensor’s BLI signal, we were able to monitor binding of 3 pM phage (Appendix C.8.b). Taking into account that Nb-displaying phage were produced by infecting bacteria containing a pIII phagemid display vector with helper phage, less than 10 % of our phage display one Nb on their tip (Russel et al., 2004), meaning we measured ~0.3 pM of phage only. Notably, we observed a decrease in the BLI signal upon phage binding. This phenomenon has also been observed for the binding of whole cells, where scattering results in a large change in optical thickness at the biosensor surface (Li and Wassaf, n.d.; Verzijl et al., 2017): a small increase in optical thickness shifts the interference pattern to the right compared to an internal reference, but a big increase shifts it to the left, causing signal inversion (Cameron et al., n.d.). Contrarily, the association curves of the unpurified phage contained in a culture supernatant are positive (Appendix C.8.c), possibly because the composition of the supernatant limits scattering. Within 10 min of association of 0.4 nM Nb207-phage, we observed that a GFP-sensor is saturated. However, 10 nM of the Nb207 alone is not enough to saturate the same sensor within 10 min. It thus appears that not all the binding sites on the sensor are accessible to the phage, possibly by steric hindrance due to their size.

### Enrichment of binders over panning rounds

Though we used sensors that are saturated with the target protein to exploit the full capacity of the sensors, we found that the number of phage that binds to the sensor in a first round is too low to detect association by interferometry. When panning a purified phage library on GFP, phage binding could only be monitored in the second round, causing a significant decrease upon binding and an increase back to baseline upon dissociation and elution of the remaining phage (Figure 2). Interestingly, in pannings with unpurified phage, binding results in a bigger positive signal (Figure 3), adding another advantage to panning with unpurified phage. Sensorgrams of irrelevant Nb-phage can be subtracted to correct for the (minor) baseline shift but the signal to noise was considerable and we found that differences in amplitudes amongst similar experiments correlates with the amount of phage that binds and elutes. Accordingly, the enrichment of binders can be followed over panning rounds via association curves to omit time-intensive titration steps.

### Octet pannings can be downscaled

We also found that the amount of (purified) target needed for selections can be reduced and the throughput can be increased by panning in 384-well plates. In our hands, Octet pannings on GFP in flat (80 µL) or tilted bottom (40 µL) 384-well plates resulted in the same enrichment as in 96-well plates containing 200 µL (insert in Figure 3). One downside of reducing the well volume is that less phage are added during association, resulting in a lower binding signal. For highly diverse libraries however, it might be useful to upscale experiments in a first round to exploit the full diversity of the library. This could be achieved by spreading the phage over multiple wells or increasing the phage concentration by purification.

### Octet panning is a versatile panning method for separating subsets of binders to challenging targets under well-controlled experimental conditions

One of the major advantages of Octet panning is that experimental parameters can be precisely controlled so that specific subsets of binders can be separated in a single experiment. For example, a 10 min dissociation step allowed us to recover a high diversity of lower affinity binders for GFP, but very strong binders were collected from the same experiment by eluting in a separate well after 15 h dissociation (Appendix C.9). To show that Octet panning is also compatible with more challenging targets, where the purified (recombinant) protein is not available, a selection was performed on mouse brain membrane protein IgSF8, a critical regulator of hippocampal CA3 microcircuit connectivity and function (Apóstolo et al., 2020). Because IgSF8 was not available as a purified protein, a llama was immunized with the membrane fraction of mouse brain extract enriched in synaptic vesicles. IgSF8 fused to GFP was expressed in HEK cells, solubilized in DDM and immobilized on a sensor using GFP-specific Nb207. Because we anticipated low abundance of target-specific phage, we increased the association time to 2 h and washed twice for 1 min before recovering binders in acid. After 2 rounds, 53 % of the characterized clones were antigen-specific. Next, we purified one representative of a family of medium binders (Nb776) and another from a family of strong binders (Nb777) to confirm specific binding to IgSF8-EGFP by immunoprecipitation (Appendix C.10). These experiments show that Octet panning is easily tuned to discover target-specific binders against challenging membrane proteins without the need for highly purified protein.

### Octet pannings are amenable to throughput approaches

To showcase the throughput of our method, two rounds of Octet panning were performed in parallel on five different yeast proteins using a standard 8-channel Octet device. A llama was immunized with the soluble fraction of a crude yeast lysate containing thousands of unpurified proteins. Five unpurified GFP-tagged yeast proteins were loaded on separate sensors using Nb207. Because we anticipated low abundance of target-specific phage, the dissociation time was decreased to 10 min. Figure 4 compares the association curves of phage during the first round and the second round for the five proteins. Remarkably, these curves appear to mirror the enrichment of target-specific phage for the different antigens on the different sensors. Indeed, the larger amplitudes for PDC1 and PGI1 coincide with the screening results. In this experimental setup, no binders were found for SSA1 or BMH1. The reasons for this can be poor abundance of these proteins in the yeast lysate that was used for immunization, low immunogenicity of these proteins, steric hindrance of the GFP-tag, amongst other explanations.

Throughout this study we showed that immobilizing a target on an Octet sensor and using the Octet device to transfer this solid phase to different wells of a microtiter plate allows to perform phage display selections under well-controlled regimes with passive dissociation and/or active elution. By omitting phage purification and titration steps, we can perform three rounds of Octet panning in five consecutive days (Appendix C.12). On an Octet R8 device, one can easily perform up to eight selections in parallel to compare different experimental conditions or to discover binders for different antigens. We anticipate that similar experiments can be scaled up to 64 parallel pannings on an Octet RH96 by combining 96 sensors with two 384-well plates for target loading, phage association, washes, time-controlled dissociation and elution of bound phage. In conclusion, we show that Octet selections are a full-fledged alternative to conventional pannings to discover Nanobodies. Experiments can easily be automated to perform tunable (incubation time) pannings under precisely controlled conditions (buffer composition, temperature, …) and are amenable to throughput approaches or parallel selections where one parameter is varied (target concentration).

## CRediT authorship contribution statement

P. De Keyser: Conceptualization, Methodology, Software, Investigation, Writing - Original Draft. V. Kalichuk: Resources, Writing - review & editing. T. Zögg: Resources, Writing - review & editing. A. Wohlkönig: Resources, Writing - review & editing. S. Schenck: Resources. J. Brunner: Supervision. E. Pardon: Resources, Writing - review & editing. J. Steyaert: Supervision, Writing - review & editing.

## Acknowledgements

The graphical abstract was created with BioRender.com. We thank H. De Greve for providing the mutated GFP-expressing *E. coli* strain and R. Willaert for providing the engineered yeast strains from the Yeast GFP fusion collection. We thank J. De Wit for providing the plasmid of IgSF8-EGFP and A. Lundqvist for giving a training in phage library preparation and phage titration.

We acknowledge the support and the use of resources of Instruct-ERIC, part of the European Strategy Forum on Research Infrastructures (ESFRI), and the Research Foundation - Flanders (FWO) for their support to the Nanobody discovery.

The work described in this article has been carried out in accordance with Directive 86/609/EEC for animal experiments.

## Appendix A. Implementation Details

### Appendix A: kinetic model

*A* is antigen, the target on the sensor

*N* is Nb, the phage-displayed antigen binder in the well

*AN* is Nb bound to antigen

[ ] is the volume concentration (M = mol L^−1^)

[ ]* is the surface concentration (mol m^−2^)

[ ]_0_ is the concentration at time *t* = 0

*a* is the specific surface area (m^2^ L^−1^)

*k_on_* is the association rate constant (M^−1^ s^−1^)

*k_off_* is the dissociation rate constant (s^−1^)

It was assumed the initial surface concentration of antigen 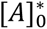 is low enough that bound phage do not overlap with free binding sites so Langmuir reaction kinetics can be used. This is a reasonable assumption given the area per binding site is minimum 280 nm^2^ and the diameter of filamentous M13 phage is 6 to 7 nm (Carmen and Jermutus, 2002):

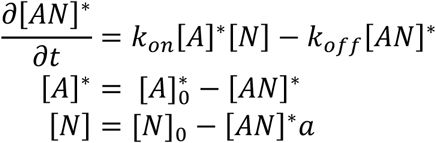

### Appendix B: sequences

GFP (CA12745):

MGHHHHHHGSKGEELFTGVVPILVELDGDVNGHKFSVSGEGEGDATYGKLTLKFICTTGKLPVP WPTLVTTLTYGVQCFSRYPDHMKRHDFFKSAMPEGYVQERTISFKDDGNYKTRAEVKFEGDTLV NRIELKGIDFKEDGNILGHKLEYNYNSHNVYITADKQKNGIKANFKIRHNIEDGSVQLADHYQQNTP IGDGPVLLPDNHYLSTQSALSKDPNEKRDHMVLLEFVTAAGITHGMDELYK

Nb207 (CA12760):

QVQLVESGGGLVQAGGSLRLSCAASGRTFSTAAMGWFRQAPGKERDFVAGIYWTVGSTYYADS AKGRFTISRDNAKNTVYLQMDSLKPEDTAVYYCAARRRGFTLAPTRANEYDYWGQGTQVTVSSH HHHHHEPEA

Nb394 (CA19394):

QVQLVESGGGLVQAGGSLRLSCAASGRTFSVSNMGWFRQAPGKERVFVAAIGWTTGSTYYADS VKGRFTISRDNTKNTVYLQMNSLKPEDTAVYRCAARRRGYSRVPMTPDEYEYWGQGTQVTVSS HHHHHHEPEA

Nb395 (CA19395):

QVQLVESGGGLVQTGDSLRLSCAASLRTFKSYAMGWFRQAPGKEREAVATISAEGDSTYYDDSV KGRFTISRDNAANTVYLQMNSLKPEDTAVYYCAASRFPYGLRLLQRRESDYWGQGTQVTVSSHH HHHHEPEA

Nb776 (CA19776): QVQLVESGGGLVQPGGSLRLSCAASGFTFTNYWMGWGRQAPGKGLEWVSAIAPGEGSTYYAD SVKGRFTTSRDNAKNTLYLQMNSLKPEDTAVYYCARIIRVVRGGRPYESDYRGQGTQVTVSSHH HHHHEPEA

Nb777 (CA19777):

QVQLVESGGGLVQAGGSLRLSCAASERTFSRNAVGWFRQAPGKEREFVATIRWSSGITYYSDSV KGRFTISKDNAKNTAYLQMNKLKPEDTAVYYCAASPVFITTRGSDYDNWGQGTQVTVSSHHHHH HEPEA

### Appendix C: tables and figures

**Appendix C.1:**
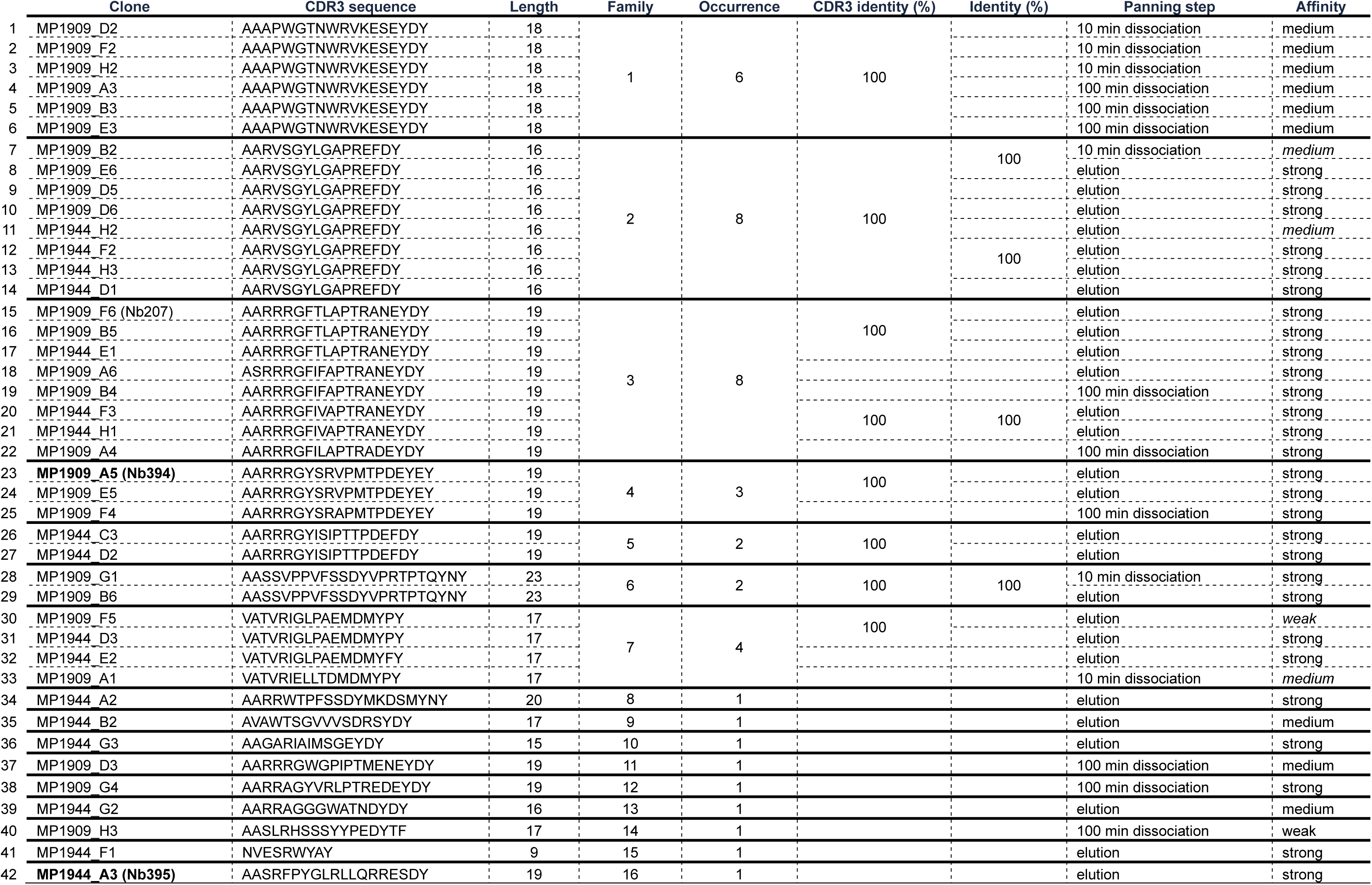
CDR3 alignment of GFP binders. Sequences of binders screened from 2 Octet pannings on GFP and Nb207 are shown. Sequences were aligned in CLC Workbench 22. Clones are family if the CDR3 has the same length and > 80 % identity. Discrepancies between the affinity of a clone determined via BLI compared to most other clones in the family are *cursive*. Clones selected for further characterization are **bold**.

**Appendix C.2:**
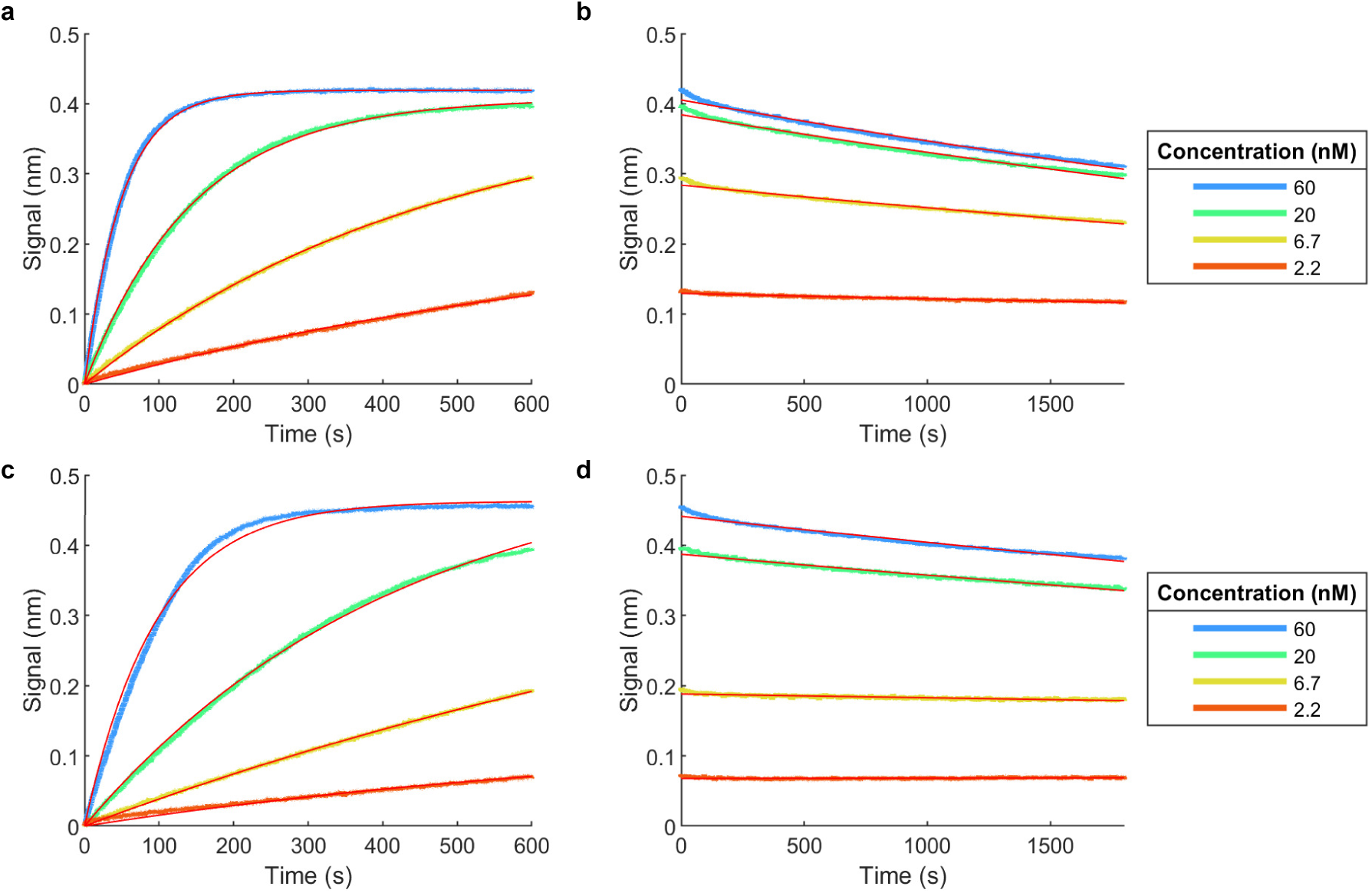
Octet characterization of 2 GFP binders selected with Octet panning. **a**. association curves of Nb394, **b**. dissociation curves of Nb394, **c**. association curves of Nb395, **d**. dissociation curves of Nb395. Data analysis was done in MATLAB, first smoothing the data, removing the baseline shift at the start of dissociation and aligning all curves at the start of association. Red lines show a one phase decay model fitted through the dissociation curve (“GraphPad Prism 9 Curve Fitting Guide - Equation: One phase decay,” n.d.) and an association kinetics model fitted through the association curve (“GraphPad Prism 9 Curve Fitting Guide - Equation: Association kinetics (two ligand concentrations),” n.d.). R^2^ is 0.9987, 0.9989, 0.9996, 0.9962 in a, 0.9876, 0.9862, 0.9802, 0.8867 in b, 0.9924, 0.9986, 0.9990, 0.9699 in c, 0.9733, 0.9716, 0.7142, 0.0416 in d.

**Appendix C.3:**
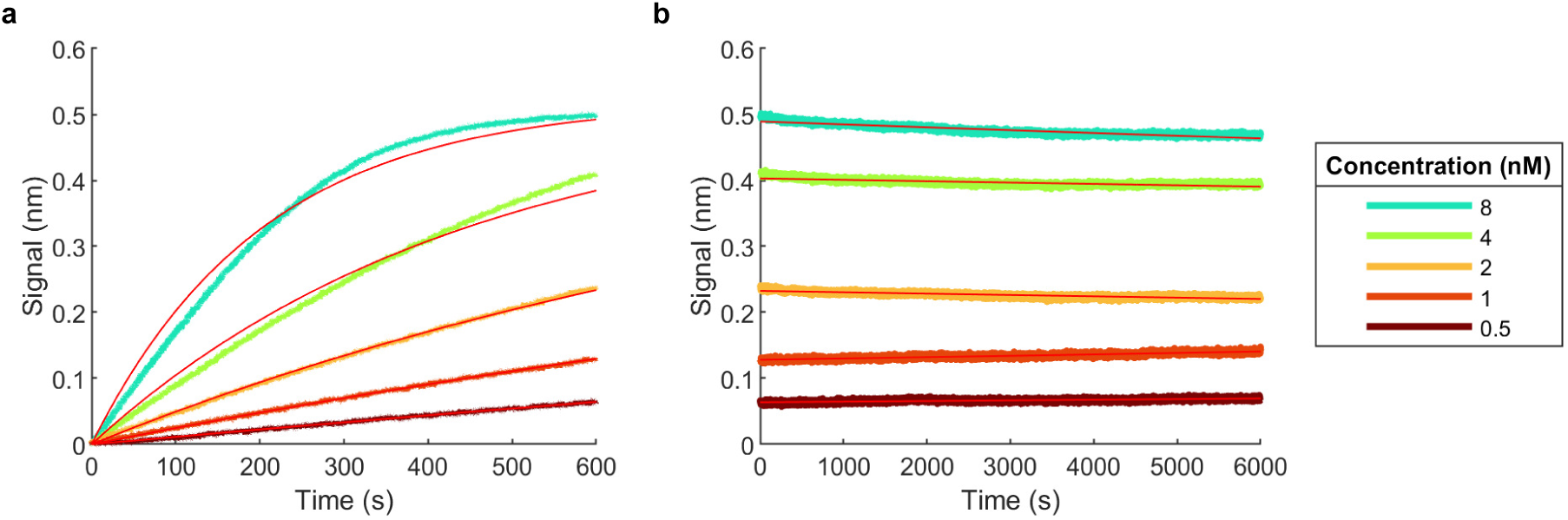
Octet characterization of GFP binder Nb207 selected previously (Steyaert et al., 2021). **a**. association curves of Nb207, **b**. dissociation curves of Nb207. Data analysis was done in MATLAB, first smoothing the data, subtracting a reference curve (sensor incubated with buffer instead of Nb207) and aligning all curves at the start of association. Red lines show a one phase decay model fitted through the dissociation curve (“GraphPad Prism 9 Curve Fitting Guide - Equation: One phase decay,” n.d.) and an association kinetics model fitted through the association curve (“GraphPad Prism 9 Curve Fitting Guide - Equation: Association kinetics (two ligand concentrations),” n.d.). R^2^ is 0.9845, 0.9871, 0.9989, 0.9981, 0.9938 in a, 0.8505, 0.6102, 0.7277, 0.8127, 0.5042 in b.

**Appendix C.4:**
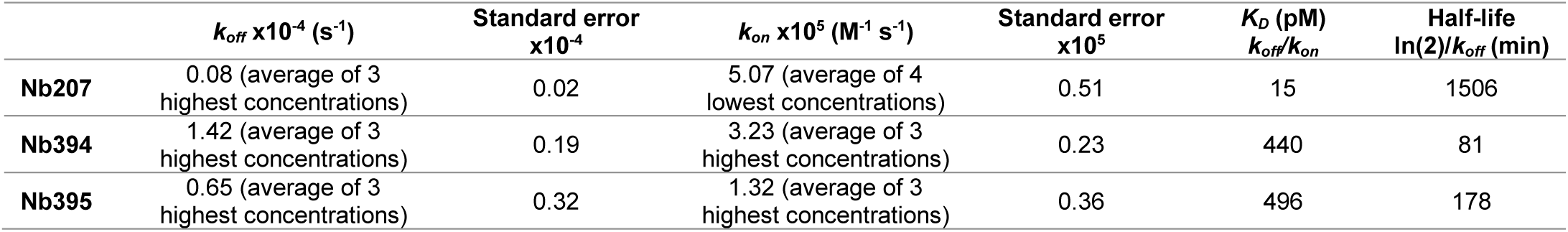
Kinetic constants of Nb207 and 2 GFP binders selected with Octet panning.

**Appendix C.5:** Number of phage eluted with different elution methods.

| Elution | Duration (min) | Phage eluted x10<sup>5</sup> |
| --- | --- | --- |
| 200 mM glycine pH 2.3 | 1 | 50 |
| 200 mM glycine pH 2.3 | 20 | 70 |
| 0.25 mg/mL trypsin | 20 | 7 |
| PBS | 20 | 3 |

**Appendix C.6:**
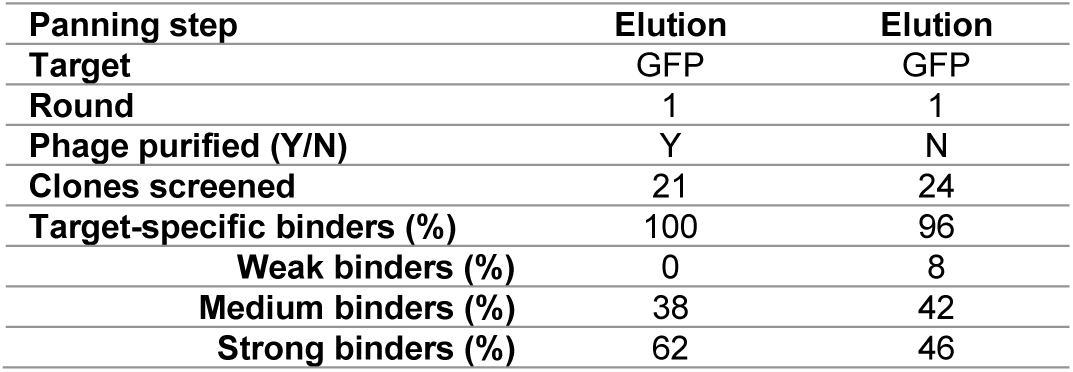
Classification of screened clones of Octet panning with purified and unpurified phage. We used Octet to present GFP that was loaded on a streptavidin functionalized sensor to a Nb library displayed on phage derived from a llama that was immunized with GFP. Classification of the GFP-specific binders resulting from a panning with purified phage and a panning with unpurified phage contained in the supernatant of a bacterial culture are shown: weak binders (half-life < 1 min), medium binders (1 min ≤ half-life < 10 min) and strong binders (10 min ≤ half-life).

**Appendix C.7:** Number of phage eluted in Octet panning with purified and unpurified phage.

| Phage eluted x10<sup>5</sup> | Purified phage | Unpurified phage |
| --- | --- | --- |
| Round 1 | 1.5 | 0.38 |
| Round 2 | 300 | 76 |
| Phage eluted round 2/round 1 | 200 | 200 |

**Appendix C.8:**
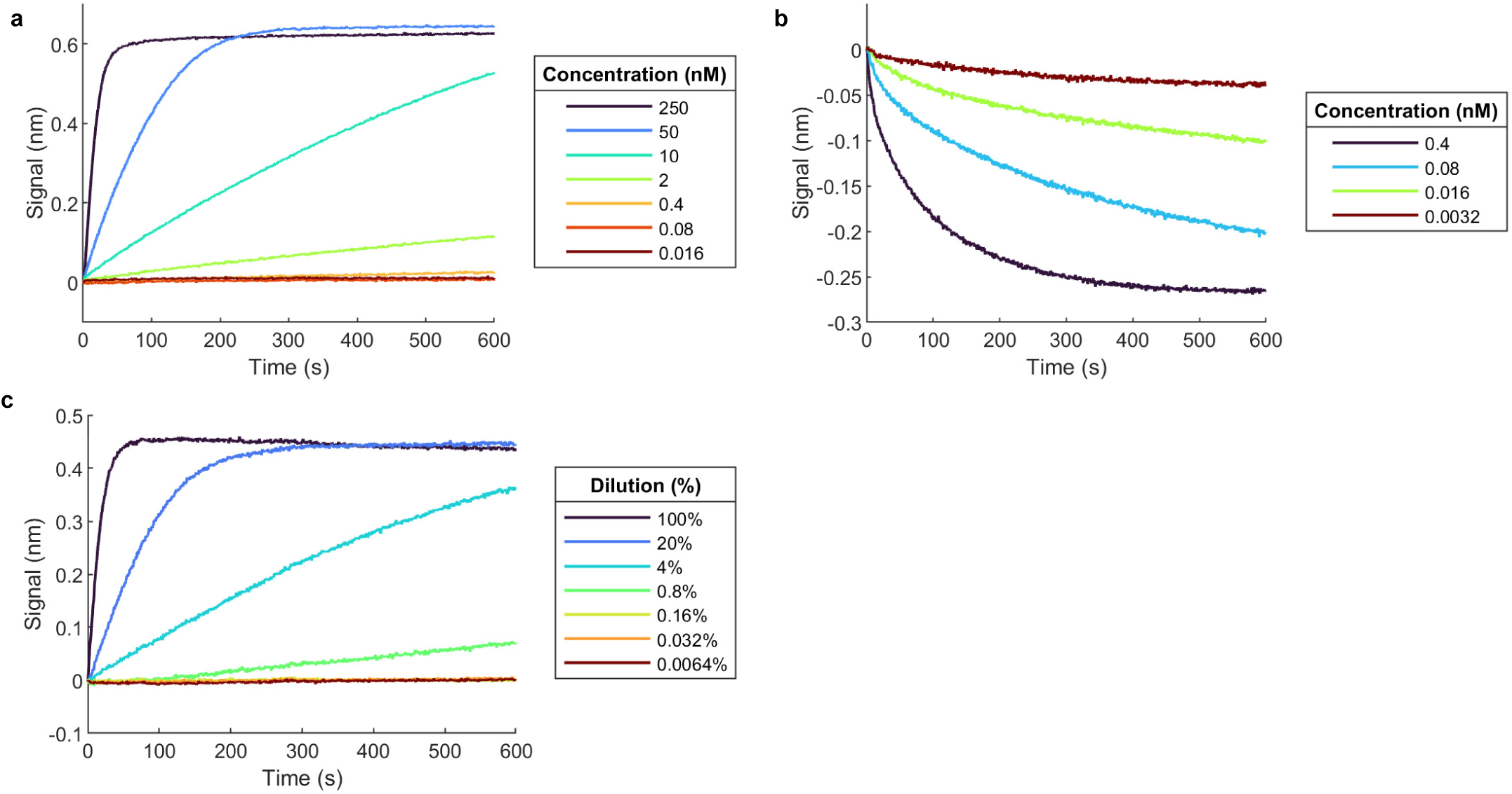
Octet association curves. Biotinylated GFP was loaded on streptavidin sensors and dipped in PBS/Tween/BSA (PBSTB) with different concentrations of **a**. Nb207, **b**. purified Nb207-phage, **c**. unpurified Nb207-phage culture supernatant for 10 min. Data analysis was done in MATLAB, first smoothing the data, subtracting a reference curve (sensor incubated with PBSTB instead of Nb207 or Nb207-phage) and subtracting the effect of culture supernatant (sensor loaded with an irrelevant protein) before aligning all curves at the start of association.

**Appendix C.9:** Classification of screened clones of Octet panning with GFP and unpurified phage. We used Octet to present GFP that was loaded on a streptavidin functionalized sensor to a Nb library displayed on phage derived from a llama that was immunized with GFP. Classification of the GFP-specific binders resulting from a panning with unpurified phage contained in the supernatant of a bacterial culture and 15 h dissociation are shown: weak binders (half-life < 1 min), medium binders (1 min ≤ half-life < 10 min) and strong binders (10 min ≤ half-life) of which very strong binders (80 % or more bound after 10 min).

| Panning step | 3 min wash | 10 min dissociation | 100 min dissociation | 500 min dissociation | 900 min dissociation | Elution |
| --- | --- | --- | --- | --- | --- | --- |
| Target | GFP | GFP | GFP | GFP | GFP | GFP |
| Round | 1 | 1 | 1 | 1 | 1 | 1 |
| Phage purified (Y/N) | N | N | N | N | N | N |
| Clones screened | 11 | 16 | 16 | 16 | 15 | 14 |
| Target-specific binders (%) | 0 | 31 | 63 | 69 | 67 | 79 |
| Weak binders (%) | 0 | 6 | 0 | 0 | 7 | 14 |
| Medium binders (%) | 0 | 25 | 44 | 6 | 0 | 21 |
| Strong binders (%) of which | 0 | 0 | 19 | 63 | 60 | 43 |
| very strong binders (%) | 0 | 0 | 0 | 6 | 13 | 29 |
| Very strong binders/ binders (%) | 0 | 0 | 0 | 9 | 19 | 37 |
| Diversity = families/binders (%) | \ | 100 | Not defined | Not defined | 70 | 55 |

**Appendix C.10:**
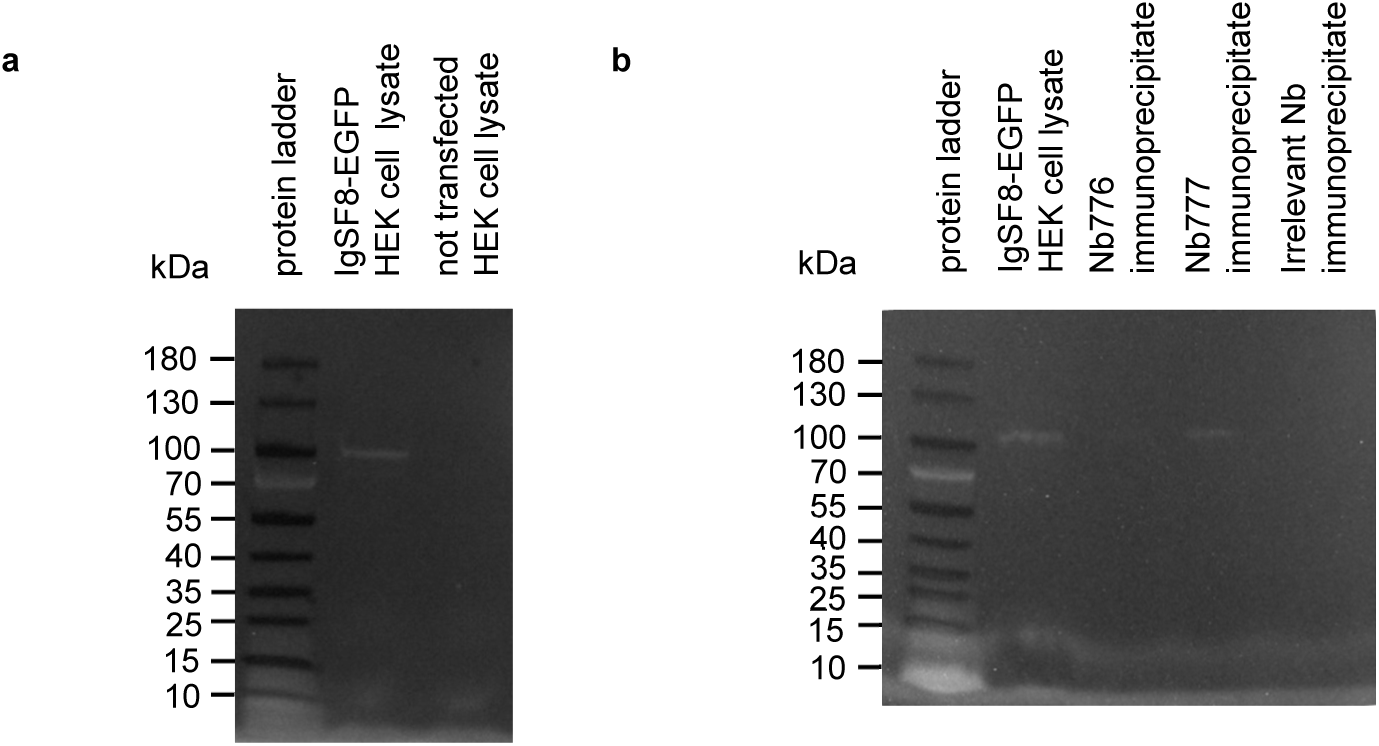
In-gel fluorescence of IgSF8-EGFP. IgSF8-EGFP expressing HEK293T cell lysate next to **a**. not transfected lysate and **b**. immunoprecipitate with Nb776, Nb777 and irrelevant Nb in 4-15 % mini-protean TGX gel. Overlay of Gel Doc EZ image on white tray (visualize ladder) and blue tray (fluorescence).

**Appendix C.11:**
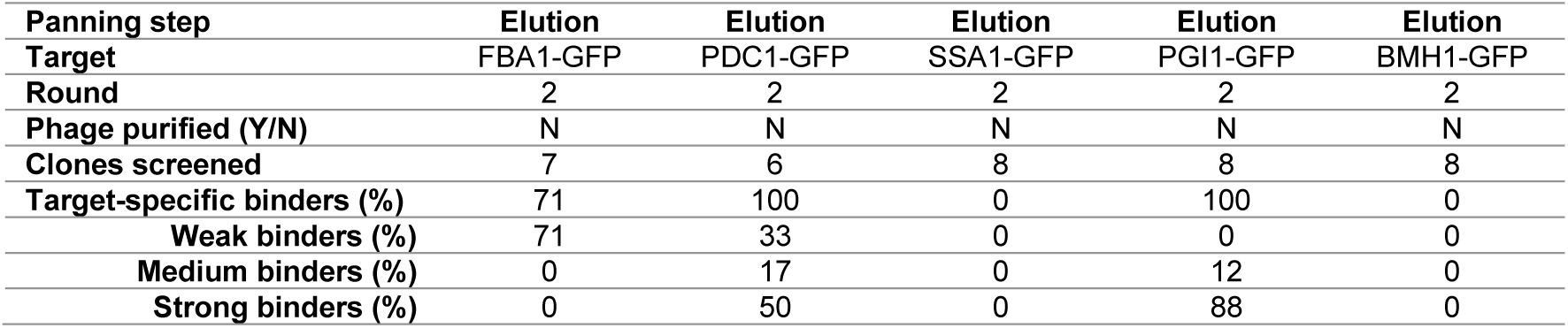
Classification of screened clones of Octet panning with five GFP-fused yeast proteins in parallel. We used Octet to present five GFP-tagged yeast proteins on Nb207-loaded sensors to a Nb library displayed on phage derived from a llama that was immunized with the soluble yeast proteome. Classification of the GFP-specific binders resulting from a panning with unpurified phage contained in the supernatant of a bacterial culture are shown: weak binders (half-life < 1 min), medium binders (1 min ≤ half-life < 10 min) and strong binders (10 min ≤ half-life).

**Appendix C.12:** 5-day schedule for 3 rounds of Octet panning.

| Step | Day | Action | Note |
| --- | --- | --- | --- |
| 1 | 1 | Grow and rescue library phage (GRL) |  |
| 2 | 2 | Round 1 with GRL |  |
| 3 | 2 | Grow and rescue elution round 1 (GR1) | When rescuing the same day, TG1 cells have not always reached OD 0.5 |
| 4 | 2 | Grow elution round 1 (G1) |  |
| 5 | 3 | Make glycerol stock from G1 |  |
| 6 | 3 | Round 2 with GR1 | Decide based on Octet signal if a 3<sup>rd</sup> round is necessary |
| 7 | 3 | Grow and rescue elution round 2 (GR2) |  |
| 8 | 3 | Grow elution round 2 (G2) |  |
| 9 | 4 | Make glycerol stock from G2 |  |
| 10 | 4 | Round 3 with GR2 | Decide based on Octet signal if a 4<sup>th</sup> round is necessary |
| 11 | 4 | Grow elution round 3 (G3) | Also rescue if a 4<sup>th</sup> round is necessary |
| 12 | 5 | Make glycerol stock from G3 |  |

